# Detection of deleterious on-target effects after HDR-mediated CRISPR editing

**DOI:** 10.1101/2020.03.27.012104

**Authors:** Isabel Weisheit, Joseph Kroeger, Rainer Malik, Julien Klimmt, Dennis Crusius, Angelika Dannert, Martin Dichgans, Dominik Paquet

## Abstract

CRISPR genome editing is a promising tool for translational research but can cause undesired editing outcomes, both on-target at the edited locus and off-target at other genomic loci. We investigated the occurrence of deleterious on-target effects in human stem cells after insertion of disease-related mutations by homology-directed repair (HDR). We identified large, mono-allelic genomic deletions and loss-of-heterozygosity that escaped standard quality controls in up to 40% of edited clones. To reliably detect such events, we developed simple, low-cost and universally applicable quantitative genotyping PCR (qgPCR) as well as sequencing-based tools and suggest their usage as additional quality controls after editing. This will help to ensure the integrity of edited loci and increase the reliability of CRISPR editing.

## Introduction

CRISPR genome editing holds great promise for biomedical research, as it allows precise and efficient genomic modifications to investigate disease-associated variants, e.g. in disease-relevant human cell types derived from induced pluripotent stem cells (iPSCs)^1,2^. However, application of CRISPR can be hampered by unwanted off- and on-target effects^3,4^. Recent studies in mice have described frequent occurrences of large deletions and complex rearrangements at CRISPR-edited loci after repair by non-homologous end joining (NHEJ)^5-8^. It is currently unclear if such alterations also affect clinically relevant human cells such as iPSCs, because repair pathways involved in CRISPR editing are differentially regulated^9^, as indicated for example by shorter human gene conversion tracts^2^. One report identified on-target effects (OnTE) at a single locus in an immortalized human cell line edited using stable overexpression of Cas9 and a guide RNA (gRNA)^6^, but did not address effects of transient expression of CRISPR machinery currently used in most editing protocols. Importantly, it has not been investigated if deleterious OnTE also occur in cells edited by homology-directed repair (HDR) to introduce specific base changes, which has high relevance in disease research, gene and cell replacement therapies. HDR- and NHEJ-edited clones are usually identified by PCR amplification of a few hundred bases around the edited locus followed by Sanger sequencing^10^, but such genotyping will fail to identify clones with large, mono-allelic insertions or deletions overlapping with genotyping primer binding sites. Instead, because the alterations prevent amplification of the affected allele, such hemizygous clones will appear to be homozygously edited (Fig. 1a). Even though false identification of homozygously edited clones can corrupt the reliability of entire studies, tests for such deleterious OnTE are still lacking in the vast majority of genome editing studies. Some reports have applied primer-walk PCR^5-8^, PacBio or other deep sequencing methods^5-7^, or droplet digital PCR^7^ to detect large on-target alterations, but these methods are expensive, laborious or require specific expertise and equipment. Here, we investigated whether large, mono-allelic deletions or insertions occur in human iPSCs after HDR-mediated CRISPR genome editing and developed quantitative genotyping PCR (qgPCR) as a simple and universally applicable tool for their reliable identification. Strikingly, we identify OnTE in up to 40% of iPSC clones edited via HDR with CRISPR/Cas9 at different loci and demonstrate deleterious effects on phenotype formation in an Alzheimer’s disease iPSC line. Extending on an earlier study^11^, we also describe large regions of copy-neutral loss-of-heterozygosity (LOH) upon HDR-mediated editing at lower frequency and validate Sanger sequencing and microarray-based tools for LOH detection. Lastly, we investigated OnTE occurrence after NHEJ-mediated CRISPR editing using qgPCR and found loss of one allele in 50% of clones.

**Figure 1.**
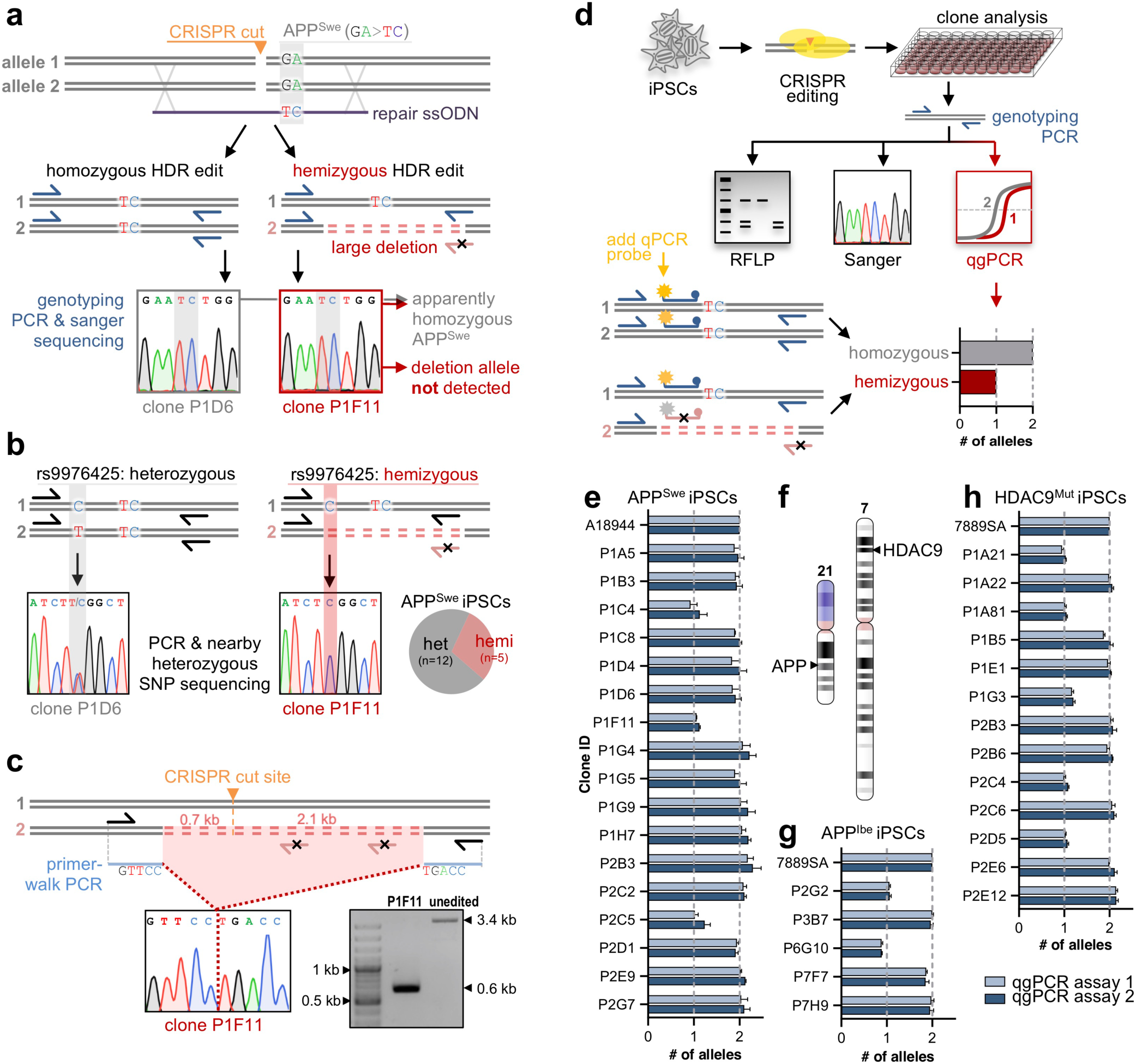
Deleterious OnTE after HDR-mediated genome editing in human iPSCs. (**a**) Sanger genotyping fails to identify mono-allelic deletions in APP^Swe^ knock-in clones. (**b**) Hemizygous APP^Swe^ clones can be detected by extending genotyping PCRs to nearby heterozygous SNP rs9976425. (**c**) Primer-walk PCR identified a 2.8 kb deletion in APP^Swe^ clone P1F11. (**d**) Adding a qPCR probe to an existing genotyping PCR allows detection of reduced allele copy numbers by qgPCR. (**e**) Allele copy numbers for two independent qgPCR assays reveal hemizygous clones with loss of one allele after HDR knock-in of APP^Swe^. (**f**) Editing positions on chromosomes 21 and 7 at APP and HDAC9 loci. (**g-h**) Identification of hemizygous clones edited at APP^Ibe^ (**g**) and HDAC9^Mut^ (**h**) loci. All values were normalized to unedited parent cell line (A18944 or 7889SA).

## Results

### Analysis of OnTE in HDR-edited iPSC clones by SNP genotyping and PCR primer-walking yields inconsistent results

To explore the incidence of deleterious OnTE in CRISPR-edited iPSCs, we analyzed 17 clones with an apparently homozygous knock-in of the APP Swedish (APP^Swe^) mutation^2^ (Fig. 1a), which causes early-onset Alzheimer’s disease. We reasoned that large, mono-allelic alterations could be identified by genotyping single nucleotide polymorphisms (SNPs) near the target site that we identified to be heterozygous before editing. Large deletions or insertions in this region would prevent amplification of the aberrant allele in a PCR covering both the target and SNP site, leading to homozygosity of the SNP in Sanger sequencing. Indeed, SNP rs9976425 appeared homozygous in 5 of 17 clones after editing, suggesting previously undetected mono-allelic changes (Fig. 1b and Supplementary Table 1). To identify possible deletions, we performed primer-walk PCRs up to 8 kb around APP^Swe^ and increased PCR extension times to detect insertions. We identified additional products in 2 of the 5 clones, revealing a deletion of 2.8 kb in clone P1F11 (Fig. 1c) and an insertion of 4.1 kb in clone P1C4 (Supplementary Fig. 1 and Supplementary Table 1). Primer-walk PCRs were, however, not able to resolve alterations in the remaining three clones identified in the SNP assay, potentially due to PCR size limitations, illustrating the requirement for more reliable readouts. In addition, both SNP genotyping and primer-walk PCRs are not universally applicable, as other loci may lack nearby heterozygous SNPs or contain regions difficult to amplify by PCR.

### Quantitative genotyping PCR (qgPCR) reliably detects widespread occurrence of OnTE in HDR-edited iPSCs

An optimal assay should not only reliably identify deleterious OnTE, but work on every edited locus, integrate well into existing gene editing workflows, and be broadly applicable with low requirements for special knowledge and equipment. As genome editing workflows usually contain a PCR for genotyping by RFLP and Sanger sequencing^10^, we reasoned that the simplest way of testing for mono-allelic alterations would be to determine allele copy number using the genotyping PCR. We addressed this by adding a labeled probe to the existing genotyping primers and performing quantitative genotyping PCR (qgPCR). Edited single-cell clones with large deletions or insertions will have higher Ct values, corresponding to a reduced allele copy number at the target site (Fig. 1d, see design parameters in Supplementary Fig. 3). To test this approach, we analyzed all 17 APP^Swe^ clones by qgPCR and confirmed the results with a second, independent qgPCR assay. Compared to unedited parent cells, three clones showed copy numbers corresponding to only one allele, which all had been previously identified by SNP genotyping (Fig. 1e and Supplementary Table 1). Interestingly, two other clones with SNP homozygosity had normal allele numbers in both qgPCR assays, suggesting LOH (confirmed in further analysis below). To investigate whether OnTE occur independently of gRNA, locus, chromosome, coding region and cell line we repeated the analysis in a different iPSC line (7889SA^2^), edited with a different gRNA for the APP Iberian mutation (APP^Ibe^). We also analyzed a line edited in a non-coding region near HDAC9 at rs2107595 (Fig. 1f), a lead SNP identified in recent GWAS studies for stroke and coronary artery disease^12^. qgPCR analysis revealed frequent loss of alleles at both loci and in both cell lines affecting 2 of 5 APP^Ibe^ and 5 of 13 HDAC9^Mut^ clones (Fig. 1g and 1h). Again, primer-walk PCRs failed to identify all affected clones. Similar to the APP^Swe^ results, SNP genotyping revealed additional clones with SNP homozygosity but normal copy number, suggesting LOH (Supplementary Table 1, see further analysis below). In agreement with previous studies^7,8^, large deletions were preferentially located at sites with microhomologies, suggesting involvement of the microhomology-mediated end joining pathway (MMEJ) (Supplementary Table 2). Taken together, our data show that deleterious OnTE, such as large deletions or insertions, occur in 18-40% of CRISPR-edited human iPSCs, and that these undesired editing events can be reliably identified by simple and universal qgPCR-based assays using already optimized genotyping PCRs.

### ‘Standard size’ qPCR assays fail to reliably detect all OnTE

Because our qgPCR assays had amplicon sizes of around 350 bp, we also tested assays with amplicon sizes of less than 150 bp, which is set as standard in most qPCR primer design tools. However, these were not reliable, because at least one assay for each analysed locus failed to identify all abnormal clones (P1C4 for APP^Swe^, P2G2 for APP^Ibe^, P1A21 for HDAC9; see Supplementary Table 1 and Supplementary Fig. 3 for further details). In all these cases, the edited loci appeared to have two normal alleles, even though there were insertions or deletions present. These InDels were missed because they did not directly overlap with the cut sites and therefore primers for short PCRs were still able to bind and support locus amplification. Hence, locus integrity cannot be reliably tested by ‘standard size’ qPCRs but requires our longer qgPCR design.

### OnTE affect phenotype formation in an iPSC-based model of Alzheimer’s disease

Most of the OnTE we found in our HDR-edited lines caused large changes on the genomic loci, which in many cases could result in major changes in gene expression, unless the allelic damage is compensated by the other allele. As most HDR-mediated CRISPR editing is performed to insert or correct disease-associated mutations, defective alleles may also have unintended effects on disease modeling. To investigate potential consequences of undesired OnTE on protein expression in a disease model, we differentiated APP^Swe^ iPSCs with and without mono-allelic alterations into cortical neurons and measured total APP levels, as well as secretion of the APP cleavage product Aβ (Fig. 2a). Hemizygous APP lines displayed a reduction in APP expression and Aβ secretion by about 50% (Fig. 2b-d and Supplementary Fig. 2). Such a reduction in Aβ levels may reduce pathogenic effects or even prevent formation of Alzheimer’s disease phenotypes in an affected iPSC-based disease model, thus illustrating potential negative effects of undetected OnTE on the reliability of studies using CRISPR/Cas9 editing for disease modeling.

**Figure 2.**
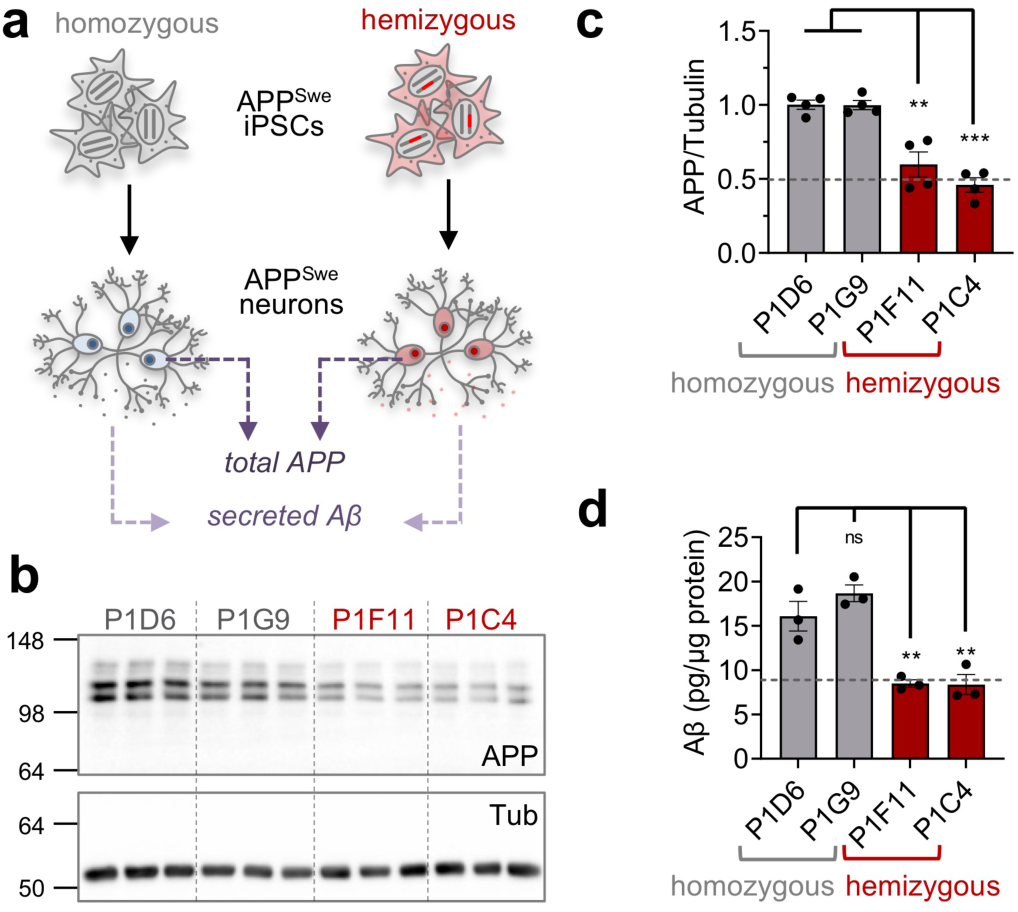
OnTE affect phenotype formation in an iPSC-based model of Alzheimer’s disease. (**a**) Two homozygous or hemizygous APP^Swe^ clones were differentiated into cortical neurons and levels of total APP and secreted Aβ were measured. (**b**) Western Blot of APP and Tubulin indicates reduced APP expression in hemizygous clones. (**c**) Quantification of (b) and biological replicates in Supplementary Figure 2 (APP normalized to Tubulin and mean of homozygous clones on same gel, n=4). (**d**) Aβ secretion (normalized to total protein amount, n=3) is also reduced in hemizygous clones. Values represent mean ± s.e.m. **P < 0.01 and ***P < 0.001, one-way ANOVA.

### CRISPR/Cas9 editing in iPSCs can cause LOH of entire chromosome arms

Our combined SNP genotyping and qgPCR analysis revealed clones with normal allelic copy number but homozygosity at nearby SNPs at all edited loci (APP^Swe^: P1G9, P2E9; APP^Ibe^: P7H9; HDAC9: P1E1, see Fig. 1e, 1g and 1h and Supplementary Table 1). We reasoned that this may result from repair of a large, mono-allelic deletion by the homologous chromosome (Fig. 3a). One previous report already indicated that copy-neutral LOH can occur after HDR-mediated CRISPR editing^11^, but it is still unclear if this is a general phenomenon or restricted to the cell line or transgene-based editing approach described in that study. To investigate the extent of LOH in our edited iPSC lines, we identified SNPs that were heterozygous in the unedited lines on both sides of the target locus up to one Mb away from the cut site and analyzed their zygosity after editing. In one clone edited at APP on chromosome 21 (P1G9) and another edited at HDAC9 on chromosome 7 (P1E1), all tested SNPs in the direction to the end of the chromosome were homozygous (Fig. 3b and 3c). Shorter regions were affected in the remaining clones (Fig. 3d). To determine whether the LOH affected the entire chromosome arm in P1G9 and P1E1 we performed whole-genome SNP genotyping using the Illumina Global Screening Array (GSA). Log R ratios showed normal copy number, but all heterozygous AB signals in B-allele frequency (BAF) were lost in the affected areas, indicating copy-neutral LOH from the cut site to the end of the targeted chromosome (Fig. 3e). Taken together, our data indicate that LOH can occur after CRISPR/Cas9 editing independently of chromosome, locus, cell line or editing method.

**Figure 3.**
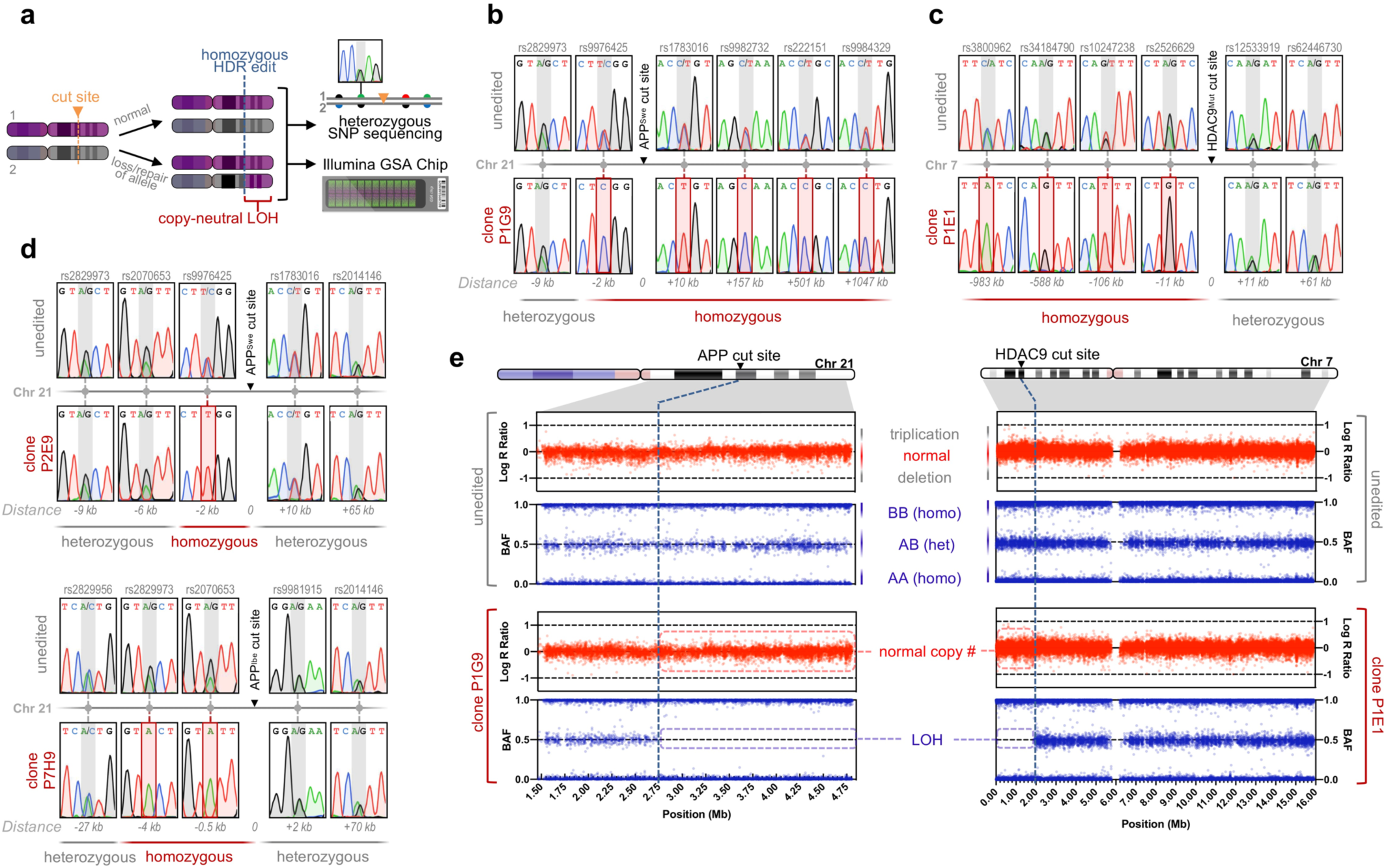
Detection of copy-neutral LOH after HDR-mediated genome editing in human iPSCs. (**a**) HDR editing may cause LOH, which can be detected via nearby SNP genotyping or SNP microarrays. (**b-c**) Sanger sequencing traces of SNPs in control and edited clones up to one Mb around APP^Swe^ (clone P1G9, **b**) or HDAC9 (clone P1E1, **c**) cut sites. (**d**) Sanger sequencing traces of SNPs in control and edited clones around APP^Swe^ (clone P2E9, top) or APP^Ibe^ (clone P7H9, bottom) cut sites. (**e**) Log R ratio and BAF in control and edited clones for chromosome 21 (P1G9 edited for APP^Swe^, left) and 7 (P1E1 edited at HDAC9, right).

### OnTE are also widespread in iPSCs edited via the NHEJ pathway

Earlier work in mice and human cell lines indicated widespread occurrence of OnTE after CRISPR editing via the NHEJ pathway^5-8^, but it is currently unclear if OnTEs are also found in iPSCs, which differ in the regulation of repair pathways^9^. We therefore analyzed APP^Swe^ clones edited via NHEJ using the same CRISPR pipeline we also applied for HDR editing. We isolated clones for which loss of a restriction site overlapping the cut site indicated presence of InDels, and further analyzed the 28 clones in which presence of two alleles could not be shown by detection of two distinct bands in gel electrophoresis after locus PCR. 12 of these clones had differently edited alleles (seen as double peaks in Sanger sequencing), indicating presences of two alleles, and this was confirmed by qgPCR in all cases. Strikingly, out of the remaining 16 clones with apparently homozygous NHEJ editing (i.e. clean single peaks in Sanger sequencing), 8 had an allele copy number of only ‘one’ in two independent qgPCR assays (Fig. 4). These results were consistent with results from our nearby SNP genotyping assay: hemizygous clones identified by qgPCR were now homozygous at SNP rs9976425 (data not shown). Thus, if researchers choose homozygously edited NHEJ clones to have a ‘clean’ knockout on both alleles, they might run into a 50% risk of using a clone with OnTE.

**Figure 4.**
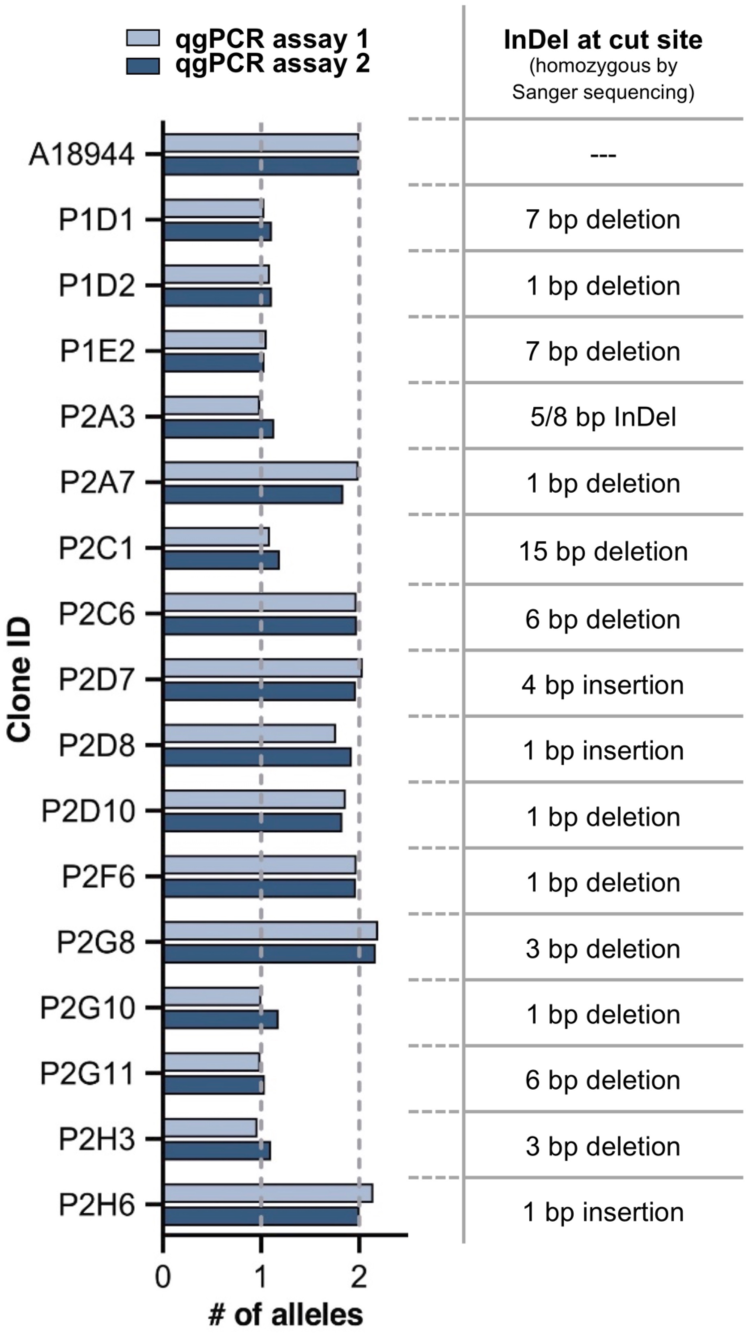
Widespread formation of OnTE after NHEJ-mediated genome editing in human iPSCs. Allele copy numbers for two independent qgPCR assays reveal hemizygous clones with loss of one allele at the APP^Swe^ locus after NHEJ editing (left). 50% of clones with apparently homozygous editing are affected. InDel sizes as determined by Sanger sequencing (right).

## Discussion

The recent CRISPR revolution has provided researchers with powerful genome editing tools that are widely applied in basic and translational research and currently also cross barriers into therapeutic applications of CRISPR-edited cells and editing directly in patients^13^. However, CRISPR editing can cause unintended effects at the edited site and elsewhere in the genome. While off-target effects can be efficiently detected with a variety of tools, the occurrence of on-target effects (OnTE) has only been described recently in mice and an immortalized human cell line. In these studies, OnTE occurred frequently upon genome editing via the NHEJ pathway, independently of the applied CRISPR system (plasmid, RNP, mRNA)^5-8^. However, it has been unclear if OnTE also occur in clinically relevant human stem cells, or after editing by homology-directed repair (HDR), which is used to introduce specific base changes.

We show that large, mono-allelic deletions and insertions occurred in 18-40% of human iPSC clones after HDR-mediated CRISPR editing. These deleterious OnTE appeared independently of the targeted locus, gRNA, coding regions, or the edited cell line, suggesting widespread prevalence of on-target issues in iPSCs, and also in other organisms and systems. By differentiating edited iPSCs with and without such unintended alterations into cortical neurons and comparing levels of Alzheimer’s disease-relevant Aβ secretion, we demonstrate the drastic effects unnoticed genomic alterations can have on studies using CRISPR-edited cells. Confirming and extending on earlier work in other systems, we also demonstrate presence of OnTE in up to 50% of iPSCs edited via the NHEJ pathway.

Furthermore, we also observed the occurrence of copy-neutral loss-of-heterozygosity (LOH) after CRISPR editing, affecting entire chromosome arms. Similar LOH has also been described in human pre-implantation embryos edited by CRISPR to correct heterozygous mutations by interhomolog recombination^14^. However, a major difference to our study is that the LOH allele did not simply acquire the sequence of the other allele, but in addition contained the mutation introduced by the repair template used for HDR, indicating a more complex repair scenario, in which it is not obvious that one allele acquired the sequence of the other. Such loss of SNP heterozygosity may potentially alter gene expression or expose effects of recessive mutations, which could be detrimental, especially in edited human embryos and clinical applications of iPSCs.

Our findings highlight the need for technologies that reliably detect all unwanted OnTE. Standard quality controls broadly performed in the field such as genotyping or karyotyping will only detect small events restricted to genotyping amplicons, or very large chromosomal aberrations, such as megabase-sized deletions, translocations and inversions, but miss the CRISPR-induced OnTE we and others revealed^5,7,8,11,15^.

Moreover, high density SNP arrays faithfully detect only larger deletions, inversions and LOH, as their reliability increases with the number of affected SNPs. LOH affecting single SNPs may be visible, but the reliability of chip data for single SNPs is lower than for Sanger sequencing, and often depends on the detection probe, genomic location etc. copy-neutral inversions are usually invisible in chip assays.

Many of the OnTE we found were small, affecting only a couple hundred to a few thousand base pairs. Accordingly, these events overlapped with no, a single, or only a few SNPs. While such small events could be reliably detected by qgPCR (deletions, insertions, inversions) or our Sanger sequencing-based assay (LOH), they could not be faithfully detected by the standard GWAS chip technology we used. Using higher density chips would not solve this problem, as the detection is not limited by the overall number of measured SNPs on the chip, but the number of measurable affected SNPs around the edited locus.

We therefore developed and validated assays based on quantitative genotyping PCR (qgPCR), Sanger sequencing and microarrays, that allow reliable detection of OnTE in iPSCs and other systems. We selected these techniques due to their simplicity, low cost, easy integration into existing workflows, universal applicability for HDR- and NHEJ-mediated CRISPR editing in various systems, and feasibility for non-specialist labs, to allow broad dissemination and acceptance in the field. We suggest using both qgPCR and nearby or global SNP genotyping as additional quality control measures to increase the reliability of CRISPR editing (see detailed workflow in Fig. 5).

**Figure 5.**
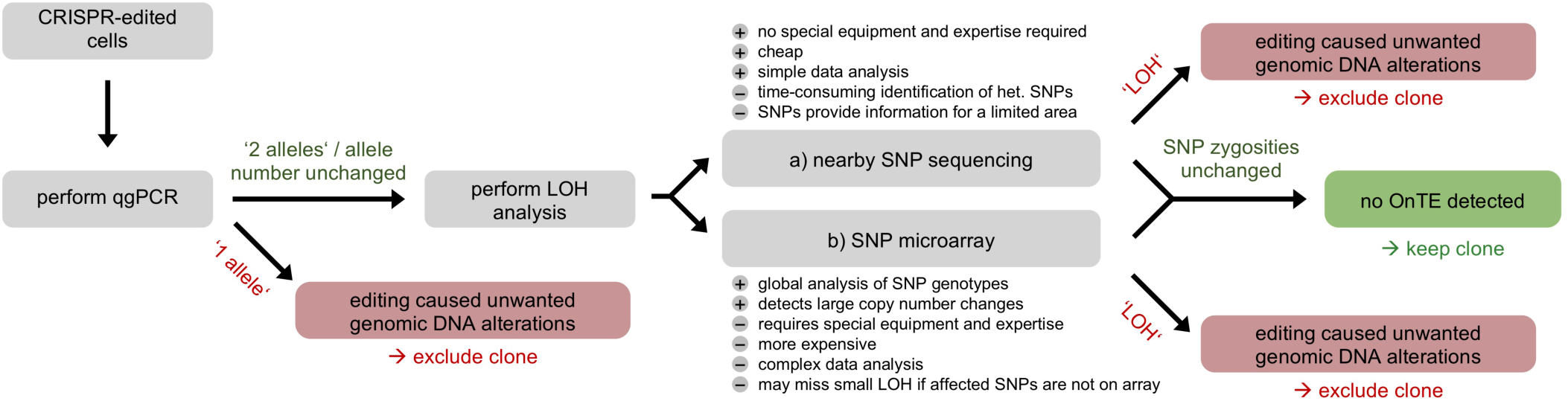
Workflow of suggested quality control experiments to determine OnTE after CRISPR editing. Single cell clones edited by CRISPR/Cas9 are first subjected to analysis by qgPCR to confirm unchanged allele numbers in edited clones and exclude clones with altered allelic copy number. To check clones for loss-of heterozygosity (LOH), there are two possibilities: nearby SNP sequencing and SNP microarrays. Both methods have their individual advantages and the selection needs to be done according to the researchers needs: While nearby SNP sequencing is cheap and does not require special equipment or expertise for analysis, SNP microarrays are more expensive and involve complex data analysis. Local SNP sequencing is more sensitive towards small regions of LOH that overlap only with few SNPs but identifying those heterozygous SNPs on both sides of the target site can be laborious in contrast to a fast analysis by microarrays. Furthermore, SNP microarrays analyze SNP genotypes genome-wide and enable characterizing the dimension of large regions of LOH, whereas nearby SNP genotyping is restricted to few loci around the edited site. Taken together, qgPCR analysis as well as nearby SNP genotyping and/or clone analysis by SNP microarrays should be conducted after CRISPR/Cas9 genome editing to ensure integrity of the edited loci.

In this study, we focused on developing reliable assays for OnTE detection to meet the urgent need of the CRISPR field for thorough quality control measures of edited cells and animals. However, future work should be aimed at not only detecting these OnTE, but understanding their biological roots and reasons for occurrence, leading to strategies to avoid their formation in the first place. This could be addressed by (1) studying locus-dependent influences, such as chromatin structure, (2) effects of genome editing reagents, e.g. by using Cas9 nickase or another nuclease, (3) effects of repair templates by modulating ssODN design and orientation, and (4) influences of other repair pathways, e.g. by modulating NHEJ or MMEJ using knockdowns or specific inhibitors.

## Acknowledgements

This work was supported by grants from the Deutsche Forschungsgemeinschaft (DFG, German Research Foundation) under Germany’s Excellence Strategy within the framework of the Munich Cluster for Systems Neurology (EXC 2145 SyNergy – ID 390857198), Vascular Dementia Research Foundation, VERUM Foundation, Wilhelm-Vaillant-Foundation, and the donors of the ADR AD2019604S, a program of the BrightFocus Foundation (to D.P.). We thank Peter Lichtner for GSA chip analysis, Johannes Trambauer and Harald Steiner for help with the APP Western Blots, Brigitte Nuscher for help with the Aβ MSD, and Benedikt Wefers for helpful comments.

## Author contributions

I.W. and D.P. designed research, I.W., J.K., R.M., J.K., D.C., A.D. performed research. I.W., J.K., R.M., J.K., M.D., D.P. analyzed and interpreted data, I.W. and D.P. wrote the manuscript with input from all authors.

---

**Supplementary Figure 1.**
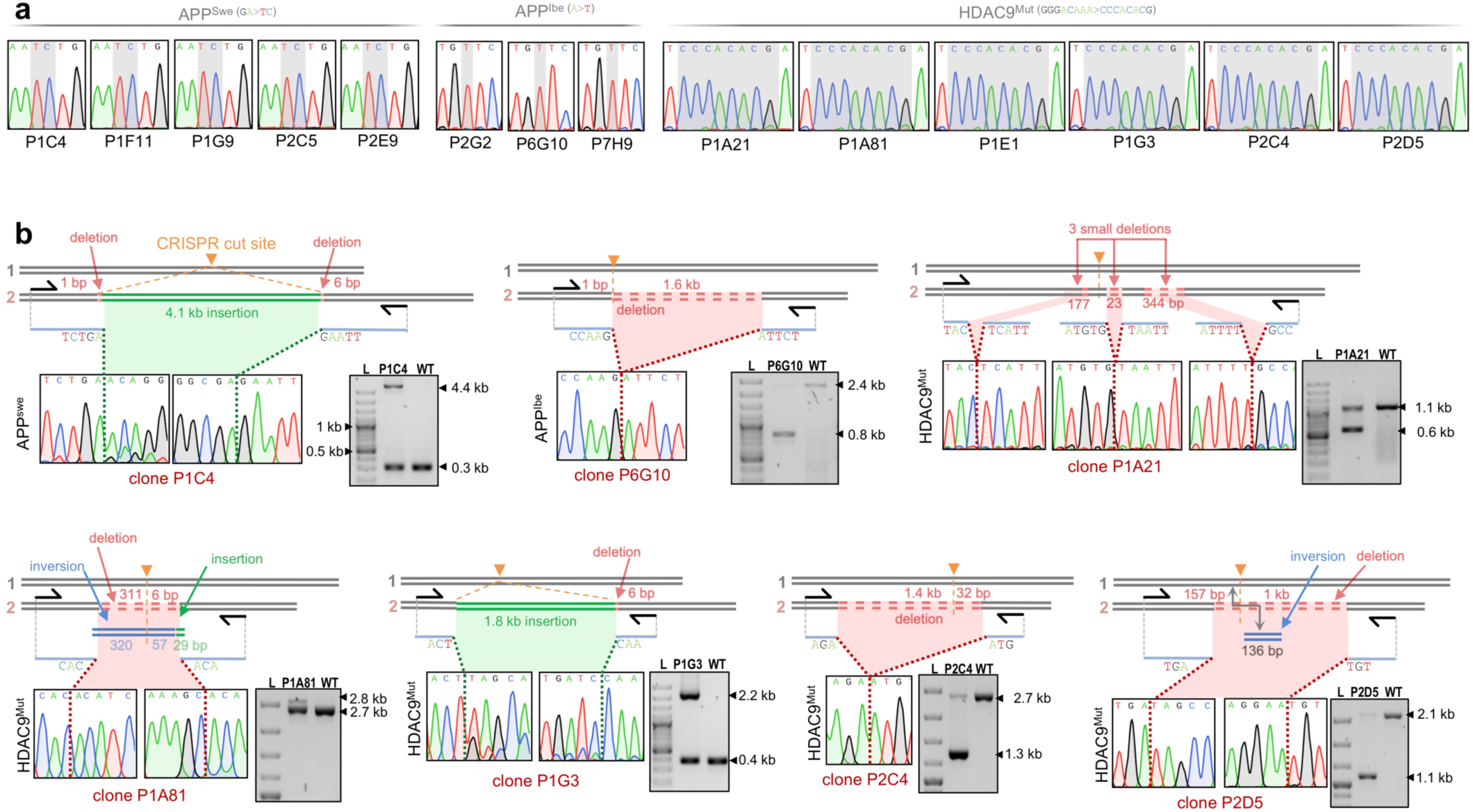
Genotyping results and primer-walk PCRs in CRISPR-edited clones. (**a**) Sanger genotyping suggests homozygous HDR editing of APP^Swe^, APP^Ibe^ and HDAC9^Mut^ in single-cell clones later identified to have mono-allelic deletions, insertions or LOH (see also Supplementary Table 1). (**b**) Primer-walk PCR identified alleles with insertions, deletions or inversions for APP^Swe^ (top left), APP^Ibe^ (top middle), and HDAC9^Mut^ clones (remaining 5 clones). L: DNA ladder, WT: wildtype. The wildtype allele is hardly visible in some clones, potentially due to preferential amplification of the shorter PCR product.

**Supplementary Figure 2.**
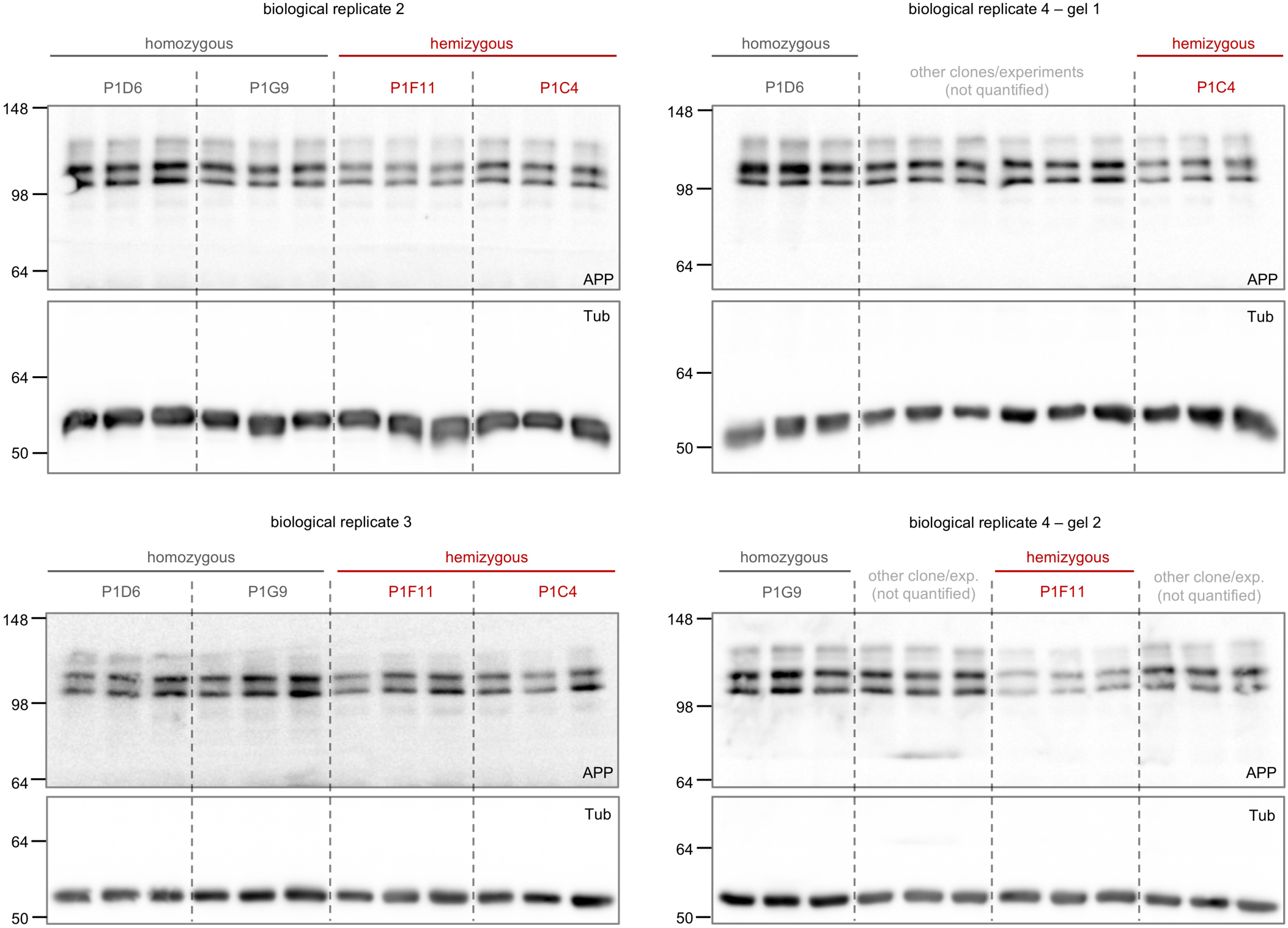
Quantification of APP expression in CRISPR-edited clones. Western blot images of biological replicates 2-4 with APP and β-Tubulin expression used for quantification shown in Figure 1l. Loading controls (β-Tubulin) were run on the same blot as APP and quantitative comparisons were only performed between samples on the same blot.

**Supplementary Figure 3.**
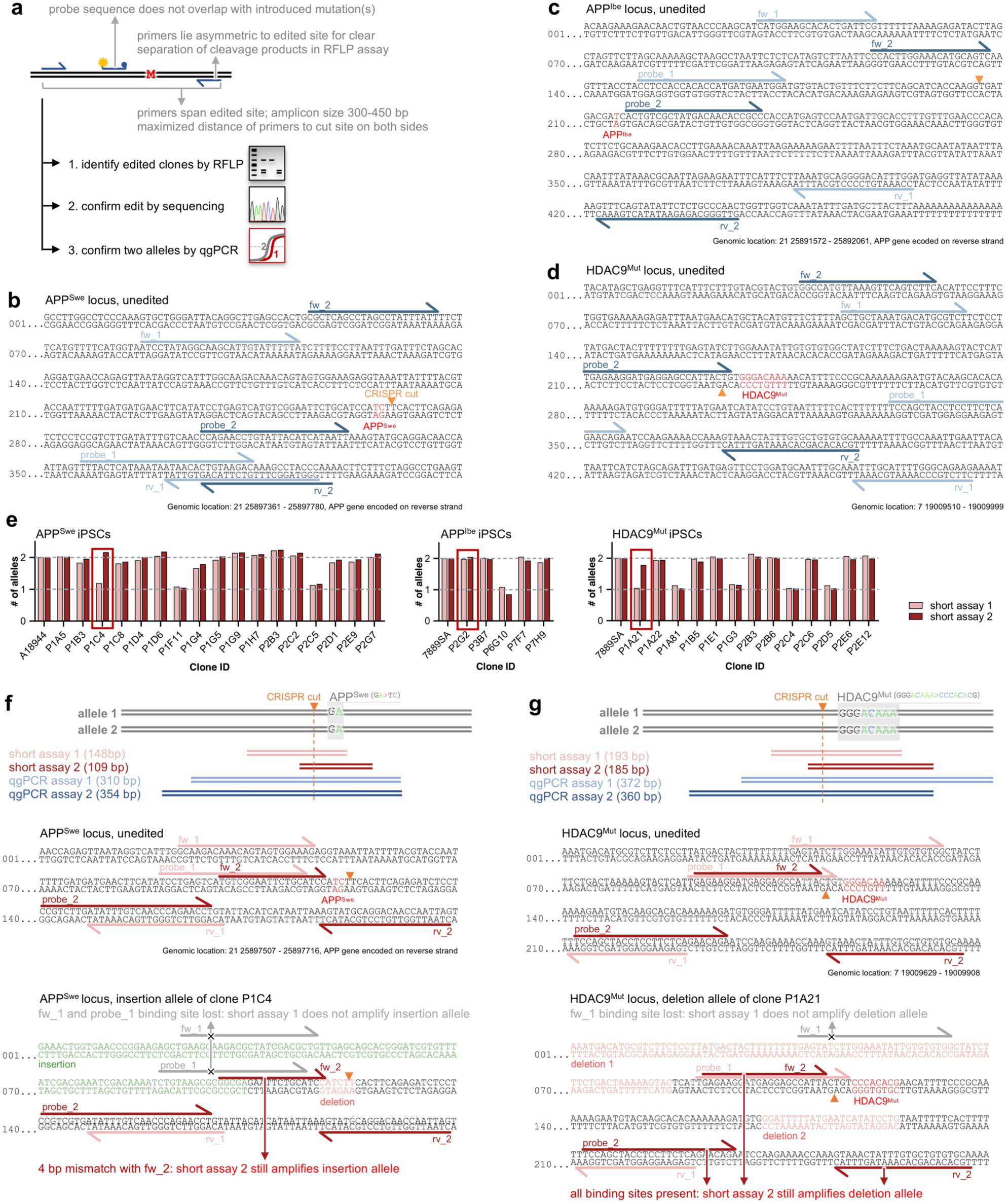
Design of quantitative PCR assays for detection of allele copy numbers in CRISPR-edited iPSCs. (**a**) Design parameters and guidelines for qgPCR assays around inserted mutation(s) ‘M’. By using the same “base PCR” as for RFLP and Sanger sequencing, qgPCR assays on edited single-cell clones easily integrate into existing genome editing workflows. (**b-d**) Positions of two independent qgPCR assays around APP^Swe^ (**b**), APP^Ibe^ (**c**) and HDAC9^Mut^ (**d**) loci shown in Figure 1, with forward primer (fw), reverse primer (rv) and qPCR probe (Full primer names, as listed in Methods: APP_Swe_Gt…, APP_Ibe_Gt…, HDAC9_Gt_…). (**e**) ‘Standard’ short amplicon qPCR assays fail to detect aberrant clones: Allele copy numbers for two independent short amplicon qPCR assays reveal several hemizygous clones, but at each locus one aberrant clone is not detected by either one or both short assays (red box). All values normalized to unedited parent cell line (A18944 or 7889SA). (**f**) Overview of qPCR probe designs (top), position of primers (fw, rv) and probes for short qPCR assays 1+2 at APP^Swe^ (full primer names, as listed in methods: APP_Swe_short…) (middle), and explanation of failed detection of insertion in clone P1C4 at APP^Swe^ locus (bottom). (**g**) Overview of qPCR probe designs (top), position of primers (fw, rw) and probes for short qPCR assays 1+2 at HDAC9 (full primer names, as listed in methods: HDAC9_short…) (middle), and explanation of failed detection of insertion in clone P1A21 at HDAC9^Mut^ locus (bottom).

**Supplementary Table 1.** Overview of CRISPR-edited iPSC clones at APP^Swe^, APP^Ibe^, HDAC9^Mut^. Data of altered clones shown in indicated figures, other data not shown.

|  |  | Copy number analysis by qPCR |  | LOH analysis |  |  | Aberrant allele |
| --- | --- | --- | --- | --- | --- | --- | --- |
| Genotype by Sanger sequencing<br>(homo-/hemizygous) | Clone ID | qgPCR assay 1+2<br>(Fig. 1e) | Short amplicon assay 1+2<br>(Suppl. Fig. 3e) | Zygosity at rs9976425*<br>(2.4 kb upstream) | Zygosity at rs1783016**<br>(10 kb downstream) | Global SNP genotyping |  |
| APP<sup>Swe</sup> | P1A5 | 2 | 2 | heterozygous | heterozygous | not tested | - |
| APP<sup>Swe</sup> | P1B3 | 2 | 2 | heterozygous | heterozygous | not tested | - |
| APP<sup>Swe</sup><br>(Suppl. Fig. 1a) | P1C4 | 1 | 1/2<br>(Suppl. Fig. 3f) | homozygous | heterozygous | no | 4.1 kb insertion, small deletions (Suppl. Fig. 1b) |
| APP<sup>Swe</sup> | P1C8 | 2 | 2 | heterozygous | heterozygous | not tested | - |
| APP<sup>Swe</sup> | P1D4 | 2 | 2 | heterozygous | heterozygous | not tested | - |
| APP<sup>Swe</sup> | P1D6 | 2 | 2 | heterozygous | heterozygous | no | - |
| APP<sup>Swe</sup><br>(Suppl. Fig. 1a) | P1F11 | 1 | 1 | homozygous<br>(Fig. 1b) | heterozygous | no | 2.8 kb deletion<br>(Fig. 1c) |
| APP<sup>Swe</sup> | P1G4 | 2 | 2 | heterozygous | heterozygous | not tested | - |
| APP<sup>Swe</sup> | P1G5 | 2 | 2 | heterozygous | heterozygous | not tested | - |
| APP<sup>Swe</sup><br>(Suppl. Fig. 1a) | P1G9 | 2 | 2 | homozygous<br>~1 Mb LOH by further SNP genotyping analysis (Fig. 3b) | homozygous | LOH until end of chromosome (Fig. 2e) | LOH |
| APP<sup>Swe</sup> | P1H7 | 2 | 2 | heterozygous | heterozygous | not tested | - |
| APP<sup>Swe</sup> | P2B3 | 2 | 2 | heterozygous | heterozygous | not tested | - |
| APP<sup>Swe</sup> | P2C2 | 2 | 2 | heterozygous | heterozygous | no | - |
| APP<sup>Swe</sup><br>(Suppl. Fig. 1a) | P2C5 | 1 | 1 | homozygous | heterozygous | no | not resolved, potentially very large deletion |
| APP<sup>Swe</sup> | P2D1 | 2 | 2 | heterozygous | heterozygous | not tested | - |
| APP<sup>Swe</sup><br>(Suppl. Fig. 1a) | P2E9 | 2 | 2 | homozygous<br>~2 kb LOH by further SNP genotyping analysis (Fig. 3d) | heterozygous | no | LOH |
| APP<sup>Swe</sup> | P2G7 | 2 | 2 | heterozygous | heterozygous | not tested | - |
| Genotype by Sanger sequencing<br>(homo-/hemizygous) | Clone ID | qgPCR assay 1+2<br>(Fig. 1g) | Short amplicon assay 1+2<br>(Suppl. Fig. 3e) | Zygosity at rs2070653*<br>(0.5 kb upstream) | Zygosity at rs5843179*<br>(0.7 kb downstream) | Global SNP genotyping | Aberrant allele |
| APP<sup>Ibe</sup><br>(Suppl. Fig. 1a) | P2G2 | 1 | 2 | homozygous | homozygous | ~ 79 kb LOH | not resolved, potentially very large deletion and LOH |
| APP<sup>Ibe</sup> | P3B7 | 2 | 2 | heterozygous | heterozygous | no | - |
| APP<sup>Ibe</sup><br>(Suppl. Fig. 1a) | P6G10 | 1 | 1 | homozygous | homozygous | not tested | 1.6 kb deletion<br>(Suppl. Fig. 1b) |
| APP<sup>Ibe</sup> | P7F7 | 2 | 2 | heterozygous | heterozygous | no | - |
| APP<sup>Ibe</sup><br>(Suppl. Fig. 1a) | P7H9 | 2 | 2 | homozygous<br>~3 kb LOH by further SNP genotyping analysis (Fig. 3d) | homozygous | ~ 10 kb LOH | LOH |
| Genotype by Sanger sequencing<br>(homo-/hemizygous) | Clone ID | qgPCR assay 1+2<br>(Fig. 1h) | Short amplicon assay 1+2<br>(Suppl. Fig. 3e) | Zygosity at rs2717369*<br>(2 kb upstream) | Zygosity at rs10260515*<br>(2 kb downstream) | Global SNP genotyping | Aberrant allele |
| HDAC9<sup>Mut</sup><br>(Suppl. Fig. 1a) | P1A21 | 1 | 1/2<br>(Suppl. Fig. 3g) | homozygous | homozygous | not tested | 3 small deletions<br>(Suppl. Fig. 1b) |
| HDAC9<sup>Mut</sup> | P1A22 | 2 | 2 | heterozygous | heterozygous | not tested | - |
| HDAC9<sup>Mut</sup><br>(Suppl. Fig. 1a) | P1A81 | 1 | 1 | 2 PCR products of different length | homozygous | no | inversion, deletion, insertion<br>(Suppl. Fig. 1b) |
| HDAC9<sup>Mut</sup> | P1B5 | 2 | 2 | heterozygous | heterozygous | no | - |
| HDAC9<sup>Mut</sup><br>(Suppl. Fig. 1a) | P1E1 | 2 | 2 | homozygous<br>~1 Mb LOH by further SNP genotyping analysis (Fig. 3c) | heterozygous | LOH until end of chromosome (Fig. 3e) | LOH |
| HDAC9<sup>Mut</sup><br>(Suppl. Fig. 1a) | P1G3 | 1 | 1 | homozygous | homozygous | no | 1.8 kb insertion, small deletion (Suppl. Fig. 1b) |
| HDAC9<sup>Mut</sup> | P2B3 | 2 | 2 | heterozygous | heterozygous | not tested | - |
| HDAC9<sup>Mut</sup> | P2B6 | 2 | 2 | heterozygous | heterozygous | not tested | - |
| HDAC9<sup>Mut</sup><br>(Suppl. Fig. 1a) | P2C4 | 1 | 1 | 2 PCR products of different length | homozygous | not tested | 1.4 kb deletion<br>(Suppl. Fig. 1b) |
| HDAC9<sup>Mut</sup> | P2C6 | 2 | 2 | heterozygous | heterozygous | not tested | - |
| HDAC9<sup>Mut</sup><br>(Suppl. Fig. 1a) | P2D5 | 1 | 1 | homozygous | homozygous | not tested | 1.2 kb deletion, inversion<br>(Suppl. Fig. 1b) |
| HDAC9<sup>Mut</sup> | P2E6 | 2 | 2 | heterozygous | heterozygous | not tested | - |
| HDAC9<sup>Mut</sup> | P2E12 | 2 | 2 | heterozygous | heterozygous | no | - |

\* PCR for SNP genotyping was spanning the cut site
\*\* PCR for SNP genotyping was **not** spanning the cut site

**Supplementary Table 2.**
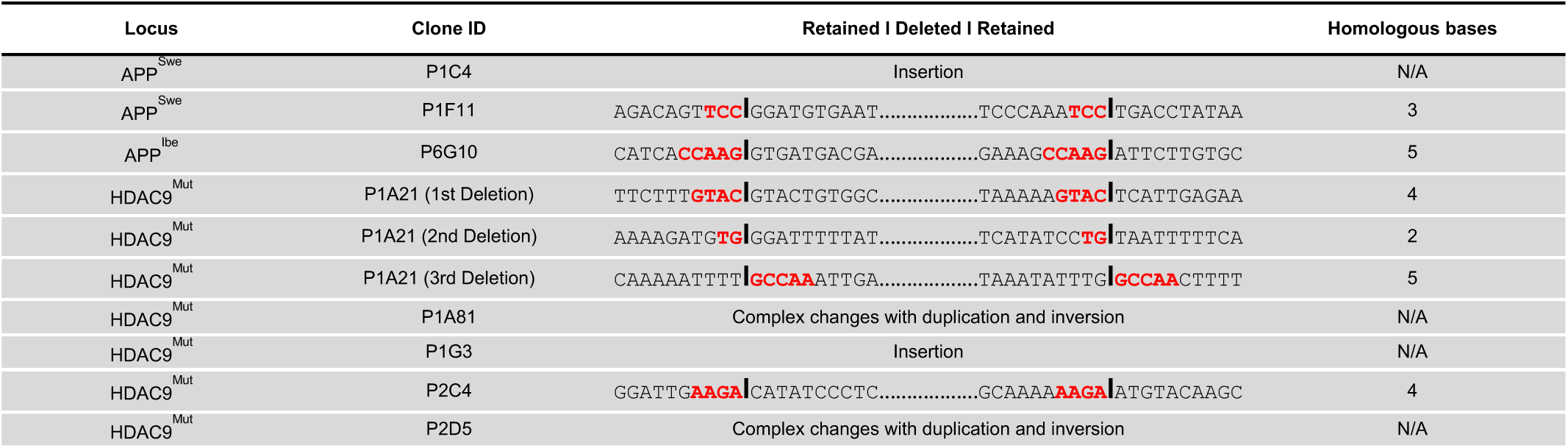
Microhomologies are prevalent at large deletion sites in CRISPR-edited iPSCs. Sequences around deletion sites with microhomologies (indicated with red letters) suggesting involvement of the MMEJ pathway. Black bars indicate sites of fusion between flanking regions.

## Methods

### Sequence information for sgRNAs, ssODNs, PCR primers and qPCR probes

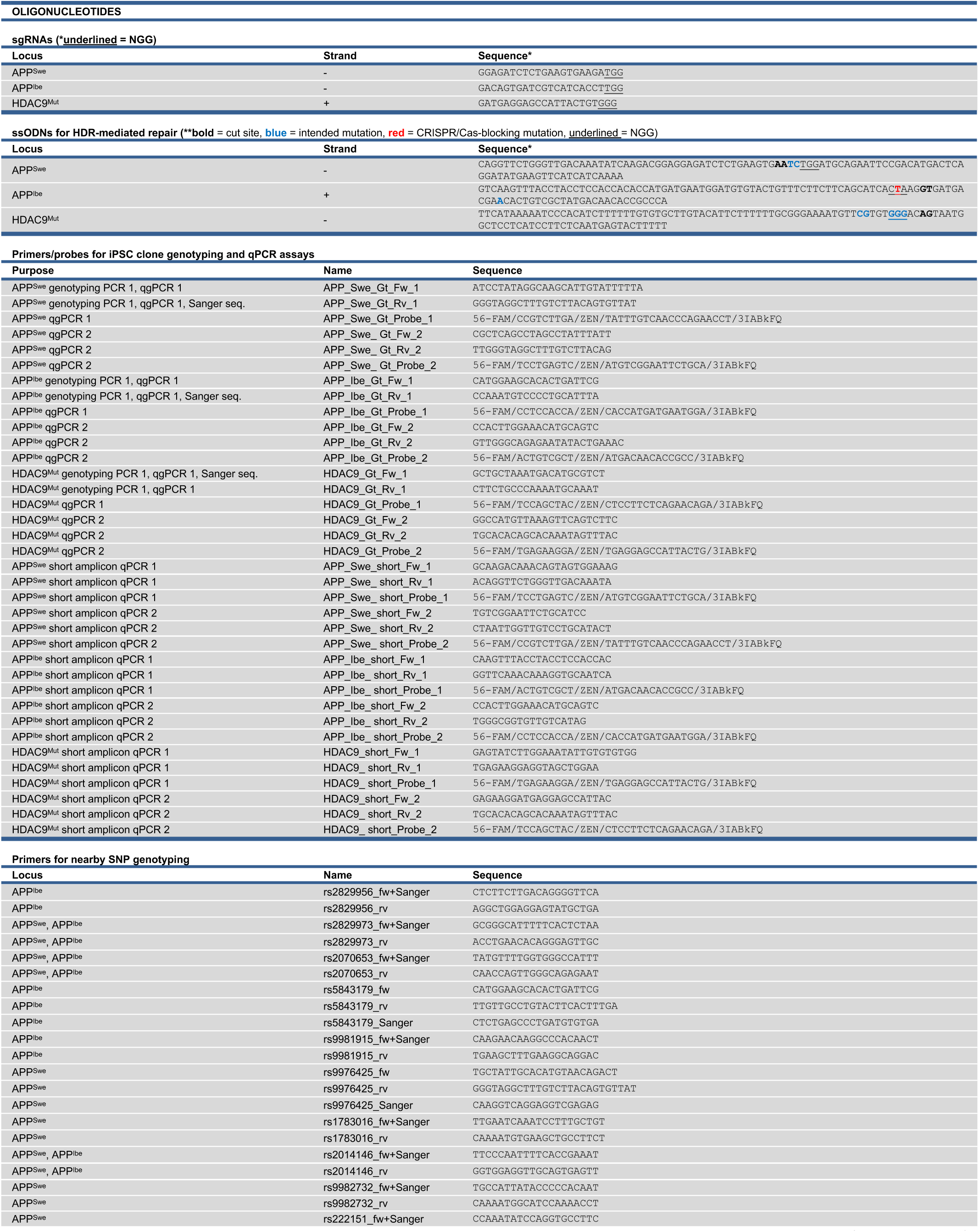

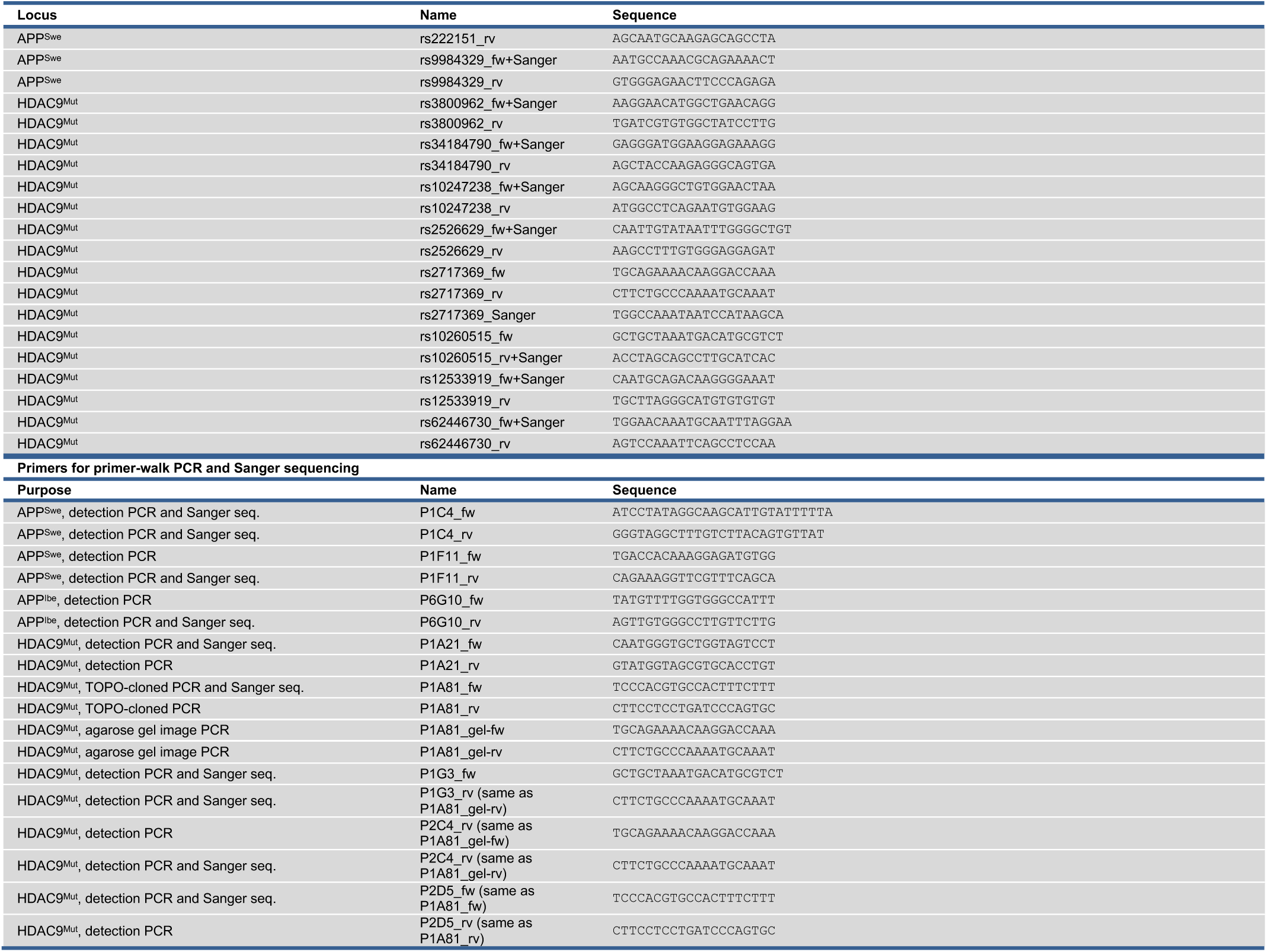

#### Lead contact and materials availability

Further information and requests for resources and reagents should be directed to and will be fulfilled by the Lead Contact, Dominik Paquet.

#### Experimental model and subject details

iPSC experiments were performed in accordance with all relevant guidelines and regulations. Work with male line 7889SA^2^ (NYSCF) was approved by the Rockefeller University Institutional Review Board after informed consent was obtained from subjects by Coriell Institute. Female iPSC line A18944 was purchased from ThermoFisher (A18945).

#### CRISPR/Cas9 genome editing

sgRNAs were designed using the CRISPR design tool (http://crispor.tefor.net/). sgRNA sequences were cloned into the BsmBI restriction site of plasmid MLM3636 (a gift from K. Joung, Addgene 43860). CRISPR editing was performed as described previously^2^ using Cas9 plasmids pSpCas9(BB)-2A-Puro (PX459) V2.0 (a gift from F. Zhang, Addgene 62988) or pCas9_GFP (a gift from K. Musunuru, Addgene 44719). Repair oligos were either symmetric 100 bp ssODNs with the same orientation as the gRNA sequence (APP^Swe,2^) or asymmetric 107 bp ssODNs (71 and 36 bp, long arm on the PAM-proximal side^16^) with sequence complementary to the gRNA (APP^Ibe^ and HDAC9), and ordered as Ultramers from IDT.

#### iPSC culture, electroporation and cortical differentiation

iPSCs were maintained on Vitronectin-coated (ThermoFisher A14700) cell culture plates and grown in Essential 8 Flex Medium (ThermoFisher A2858501) at 37°C with 5% CO2. Prior to transfection, iPSCs were transferred to Geltrex-coated (ThermoFisher A1413302) cell culture plates and grown in StemFlex Medium (ThermoFisher A3349401) containing 10 μM ROCK inhibitor (Selleckchem S1049) for two days. iPS cells were transfected by electroporation as described^10^. Briefly, two million cells were resuspended in 100 μl cold BTXpress electroporation solution (VWR 732-1285) with 20 μg Cas9, 5 μg sgRNA plasmid, and 30 μg ssODN. Cells were electroporated with 2 pulses at 65 mV for 20 ms in a 1 mm cuvette (Fisher Scientific 15437270). After electroporation, cells were transferred to Geltrex-coated 10 cm plates and grown in StemFlex Medium containing 10 μM ROCK inhibitor. Cells expressing Cas9 were selected either by sorting for GFP^10^ or selection with 350 ng/ml Puromycin dihydrochloride (VWR J593) for three consecutive days starting one day after electroporation^17^. Single-cell clone colonies were picked and analyzed by RFLP assay, using NEB enzymes TfiI for APP^Swe^, DdeI for APP^Ibe^, XmnI for HDAC9, and Sanger sequencing as previously described^10^. Cortical neuron differentiation was performed using a dual-SMAD inhibition-based protocol as described^2^.

#### Genotyping assay design and copy number analysis by quantitative PCR (qPCR)

Assays for genotyping qPCR analysis of edited single-cell clones were designed using the IDT PrimerQuest design tool. Briefly, a 400-550 bp region surrounding the edited locus was entered and the amplicon size range set to 300-450 bp. The edited site was selected as excluded region for the probe to prevent overlap. If genotyping primers were available, the primer sequences were entered under partial design input. Assays in which the probe was close to the edited site were favored. For copy number analysis, gDNA was isolated with a NucleoSpin Tissue Kit (Macherey-Nagel 740952) according to manufacturer’s instructions and 60 ng were used for analysis. As we occasionally observed variation in gDNA integrities from stored gDNA samples we recommend using fresh gDNA isolated at the same time from control and assayed clones. Freshly isolated gDNA was mixed with 2x PrimeTime Gene Expression Master Mix (IDT 1055772), 20x human TERT TaqMan™ Copy Number Reference Assay (ThermoFisher 4403316) as internal reference control, genotyping primers (0.5 pmol/μl) and the designed PrimeTime Eco Probe 5’ 6-FAM/ZEN/3’ IBFQ (0.25 pmol/μl, HPLC-purified, IDT). The quantitative PCR reaction was run for 2 min at 50°C, 10 min at 95°C, followed by 40 cycles of 15 s at 95°C and 1 min at 60°C. Allele copy numbers were determined by ddCt calculation relative to internal TERT reference and unedited control; values were multiplied by two to get total number of alleles. qgPCR experiments were performed in three independent technical replicates.

#### GSA Illumina Chip

Genomic DNA from all iPSC lines to be analysed was isolated with a NucleoSpin Tissue Kit and diluted to a concentration of 75 ng/μl. Whole-genome genotyping was performed at the Helmholtz Zentrum München (Neuherberg, Germany) using the Illumina Global Screening Array v2 genotyping chip (Illumina, San Diego, California, USA). Single nucleotide polymorphisms (SNPs) were called using the GenCall algorithm. All samples analyzed showed a sample call rate > 0.99. Gender checks were performed as an additional quality control step. SNPs with a call rate < 0.9 were discarded. All SNPs were filtered using a Hardy-Weinberg equilibrium p-value cutoff of 1E-4 and a GenTrain score cutoff of 0.7 to ensure correct clustering^18^. Log R Ratio and B Allele Frequency were extracted using Genome Studio 2.0 (Illumina, San Diego, California, USA). Data available upon request.

#### Genomic variant identification

Potential genomic variants within 5 kb around the edited loci were identified using the Ensembl Biomart tool (https://www.ensembl.org/info/data/biomart/index.html) with the following settings and filters: Ensembl variation 98 database, human Short Variants (SNPs and InDels excluding flagged variants), respective chromosome with a region of around 5kb around the edited site, global minor allele frequency >= of 0.2. The flanking sequence around the retrieved variants was downloaded from Ensembl and used in Primer3Plus (https://primer3plus.com/) to design primers for SNP genotyping. Prior to Sanger sequencing, the amplicons were analyzed for size differences by agarose gel electrophoresis to check for length polymorphisms. Heterozygosity of SNPs was confirmed by identification of double peaks in Sanger sequencing in unedited vs. edited iPSC. Heterozygous SNPs in a 1 Mb region around the edited loci were identified by parsing data from a previous molecular karyotyping experiment performed in unedited parent lines using the Illumina bead array HumanOmni2.5Exome-8 BeadChip v1.3 (Life & Brain GmbH, Bonn) (data not shown).

#### Primer-walk PCR

Primer-walk PCRs were performed with edited single-cell clones to identify aberrant PCR products with OneTaq 2x Master Mix (NEB M0486L) following manufacturer’s instructions. Primers with increasing distance to the cut site in steps of around 500 bp were tested. PCR products were analysed by agarose gel electrophoresis with a GeneRuler 100 bp Plus DNA ladder (ThermoFisher SM0321). If additional bands, not present in the unedited control cell line, were detected, PCR products were gel-purified using a NucleoSpin Gel and PCR Clean-Up Kit (Macherey-Nagel 740609) followed by Sanger sequencing. If sequencing was not successful, PCR products were TOPO cloned following manufacturer’s instructions (TOPO TA Cloning Kit for Sequencing, ThermoFisher 450030). Plasmids with TOPO-cloned inserts were isolated using the NucleoSpin Plasmid kit (Macherey Nagel 740588) and Sanger sequenced.

#### Measurements of total APP and Amyloid-β

Total protein was extracted from differentiated neurons at DIV 35 with the NucleoSpin RNA/Protein Kit (Macherey-Nagel 740933) according to manufacturer’s instructions, separated on 8% TRIS-Glycine hand-casted gels, transferred to nitrocellulose membranes (Amersham Protran 0.45 NC, GE Healthcare), boiled for 5 min in PBS, and blocked for 1 h using 0,2% I-Block (ThermoFisher T2015) with 0,1% Tween20 (Merck) in PBS. Primary antibodies (APP-Y188, Abcam ab32136, 1:4,000; Tubulin, Sigma T5168, 1:4000) were diluted in blocking solution and incubated with the membrane overnight at 4°C. After three washes in PBS + 1% Tween20, HRP-labeled secondary antibodies (Anti-Rabbit IgG (H+L), HRP Conjugate, Promega, W4011; Anti-Mouse IgG (H+L), HRP Conjugate, Promega, W4021) were added for 1h and protein signals were detected using Pierce™ ECL Western Blotting Substrate kit (ThermoFisher 32109), using a Fujifilm LAS4000 luminescence imager and band intensities quantified using ImageJ. For Aβ measurements, cell supernatant was conditioned for 5 days and experiments were performed in 3 biological replicates. Supernatants from experiments collected at different time points were flash-frozen in liquid nitrogen and stored at −80°C. Secreted Aβ1−38, Aβ1−40 and Aβ1−42 were measured with MSD Human (6E10) Aβ V-PLEX kits (Meso Scale Discovery) according to the manufacturer’s directions. Aβ values were combined to obtain total Aβ and normalized to total protein levels from cell lysate determined by Karlsson’s method^19^, as described in the NucleoSpin RNA/Protein Kit.

#### Quantification and statistical analysis

No statistical methods were used to predetermine sample size and the experiments were not randomized. Experimental data was analysed for significance using GraphPad Prism 8. Multiplicity-adjusted p < 0.05 was considered statistically significant. Significance was analysed by one-way ANOVA comparing the mean of each column with the mean of the control followed by multiple-comparison post-testing with Dunnett’s method. The analysis approaches have been justified as appropriate by previous biological studies, and all data met the criteria of the tests. The investigators were not blinded to allocation during experiments and outcome assessment.

